# The structural basis of fatty acid elongation by the ELOVL elongases

**DOI:** 10.1101/2020.11.11.378570

**Authors:** Laiyin Nie, Ashley C. W. Pike, Tomas C. Pascoa, Simon R. Bushell, Andrew Quigley, Gian Filippo Ruda, Amy Chu, Victoria Cole, David Speedman, Tiago Moreira, Leela Shrestha, Shubhashish M.M. Mukhopadhyay, Nicola A. Burgess-Brown, James D. Love, Paul E. Brennan, Elisabeth P. Carpenter

## Abstract

Very long chain fatty acids (VLCFAs) are essential building blocks for synthesis of the ceramides and sphingolipids required for nerve, skin and retina function and 3-keto acyl-CoA synthases (ELOVL elongases) perform the first step in the FA elongation cycle. Although ELOVLs are implicated in common diseases including insulin resistance, hepatic steatosis and Parkinson’s, their underlying molecular mechanisms are unknown. Here we report the structure of the human ELOVL7 elongase, which includes an inverted transmembrane barrel structure surrounding a 35 Å long tunnel containing a covalently-attached product analogue. The structure reveals the substrate binding sites in the tunnel and an active site deep in the membrane including the canonical ELOVL HxxHH sequence. This indicates a ping-pong mechanism for catalysis, involving unexpected covalent histidine adducts. The unusual substrate-binding arrangement and chemistry suggest mechanisms for selective ELOVL inhibition, relevant for diseases where VLCFAs accumulate such as X-linked adrenoleukodystrophy.

## Main

The seven human 3-keto acyl-CoA synthases (elongation of very long chain fatty acids proteins: ELOVL1-7 elongases) catalyse the first, rate-limiting step in the cycle that adds two carbon units to the acyl chains of fatty acids (FAs) with 12 or more carbons per chain (Fig. 1a). These long and very long chain FAs (LCFAs: 12C:20C and VLCFAs: >20C) ^1,2^ are the precursors for synthesis of ceramides, sphingolipids and sphingolipid signalling molecules ^3^. VLCFAs are essential for the myelin sheaths of nerves ^4,5^, the skin permeability barrier ^6,7^, retina ^8^ and liver function ^4,9^. Mutations in ELOVL elongases cause severe genetic diseases including Stargardt syndrome ^10^ and spinocerebellar ataxia ^11,12^. Mouse knockouts suggest ELOVL involvement in hepatic steatosis ^13^ and insulin resistance ^14^, and in particular ELOVL7 is implicated in cancer ^15-17^, early-onset Parkinson’s disease ^18^ and necroptosis ^19^. However, very little is known about the molecular mechanisms underlying this key step in fatty acid and lipid synthesis by the ELOVLs.

**Fig. 1.**
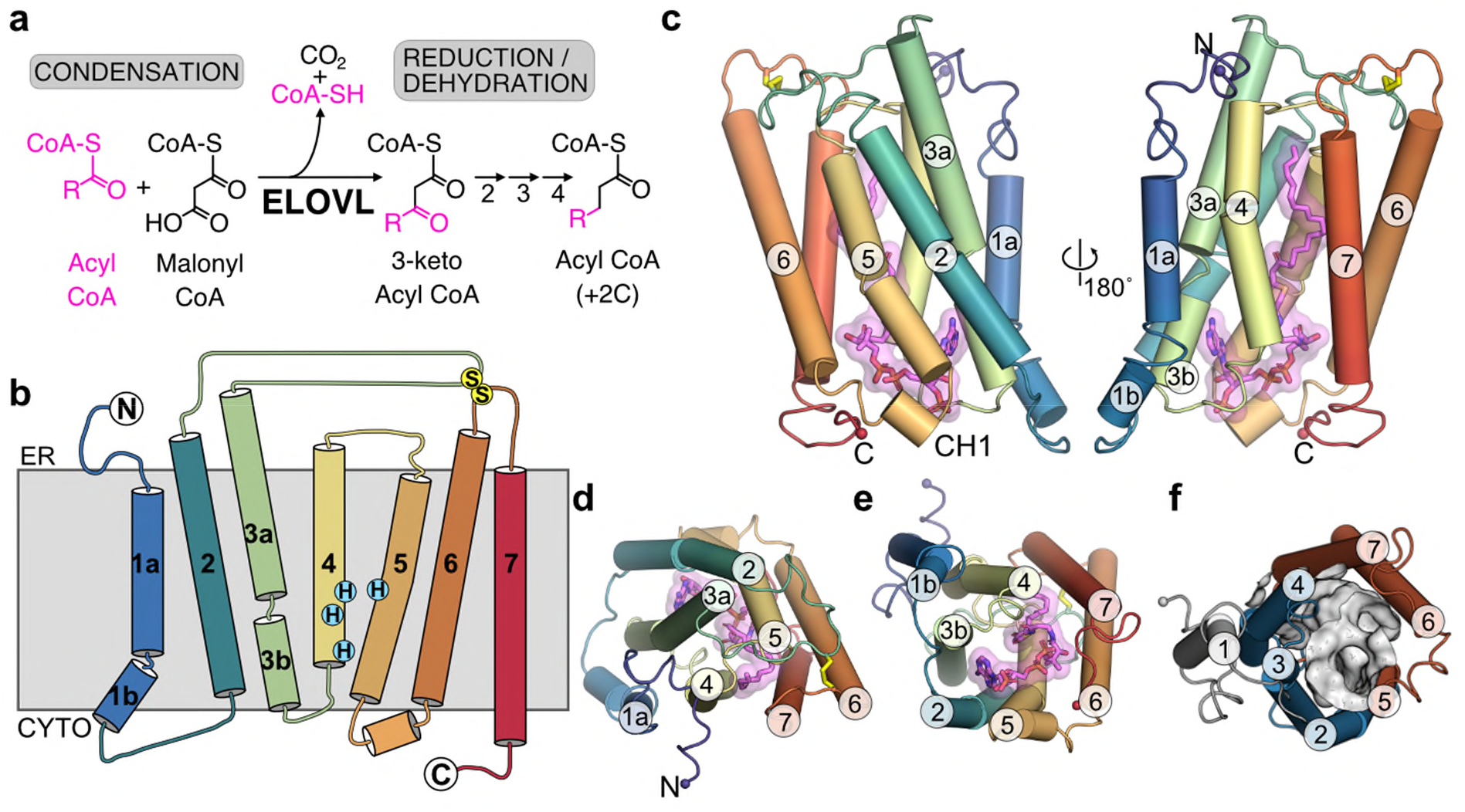
Overall structure of ELOVL7. **a**, FA elongase reaction sequence. **b**, Schematic representation of the seven TM transmembrane topology. **c-e**, Cartoon of ELOVL7 structure viewed **c**, parallel to the membrane plane and from either **d**, the ER or **e**, the cytoplasmic faces. **f**, ELOVL7 6-TM inverted barrel organisation viewed from the cytoplasmic face.

In order to understand the molecular basis of acyl chain elongation by the ELOVL elongases, we purified and solved the crystal structure of human ELOVL7 in complex with a copurified, covalently bound product analogue and performed intact protein mass spectrometry experiments to probe its catalytic mechanism. The results obtained enabled us to propose a ping-pong type mechanism, achieved through the formation of an unusual acyl-imidazole intermediate upon reaction with the first acyl-CoA substrate.

## Results

### Structure of human ELOVL7

We have solved the structure of human ELOVL7 by X-ray crystallography to 2.6 Å resolution (see Methods, Table 1 and Extended Data Fig. 1a), revealing an inverted dimer with a small and unconserved interaction surface (870 Å^2^) in the crystals which was also observed in solution (Extended Data Fig. 1b-f).

**Table 1.** X-ray data collection, refinement & validation statistics.

|  | Hg-peak | WT-native | STARANISO |
| --- | --- | --- | --- |
| **Data collection** |  |  |  |
| Space group | $P2_1$ | $P2_1$ | - |
| Cell dimensions |  |  |  |
| $a, b, c$ (Å) | 62.34, 72.42, 111.68 | 63.23, 72.46, 112.04 | - |
| $\alpha, \beta, \gamma$ (°) | 90, 100.14, 90 | 90, 100.03, 90 | - |
| Resolution limits [Å]<sup>1</sup> | 4.6, 3, 3.2 | 3.5, 2.05, 2.65 | - |
|  | (4.4, 3, 3) | (3.24, 2.05, 2.22) | - |
| **Nominal Resolution [Å]<sup>2</sup>** | **3.2** | **2.6** | - |
| Resolution [Å] | 58 - 3.0 (3.0-3.18)<sup>3</sup> | 62 - 2.05 (2.05-2.10)<sup>3</sup> | 62 - 2.05 (2.05-2.18)<sup>3</sup> |
| $R_{\text{meas}}$ | 0.396 (2.516)<sup>3</sup> | 0.161 (4.108)<sup>3</sup> | 0.119 (0.839)<sup>3</sup> |
| $R_{\text{pim}}$ | 0.093 (0.588)<sup>3</sup> | 0.061 (1.534)<sup>3</sup> | 0.062 (0.449)<sup>3</sup> |
| $I / \sigma I$ | 6.2 (1.5)<sup>3</sup> | 4.9 (0.4)<sup>3</sup> | 8.3 (1.9)<sup>3</sup> |
| $CC_{1/2}$ | 0.996 (0.838)<sup>3</sup> | 0.985 (0.473)<sup>3</sup> | 0.987 (0.813)<sup>3</sup> |
| Completeness [%] | 99.9 (99.6)<sup>3</sup> | 99.5 (96.6)<sup>3</sup> | 58.7 (17.5)<sup>3</sup> |
| Redundancy | 17.6 (18)<sup>3</sup> | 6.9 (6.9)<sup>3</sup> | 6.9 (6.2)<sup>3</sup> |
| $CC_{\text{anom}}$ | 0.251 (0.0)<sup>3</sup> | | |
| **Refinement** |  |  |  |
| Resolution (Å) |  | - | 28.68 – 2.05 |
| No. reflections (free) |  | - | 36690 (1765) |
| $R_{\text{work}} / R_{\text{free}}$ | | - | 21.04 / 22.61 |
| No. atoms |  |  |  |
| Protein |  | - | 4260 |
| Ligand |  | - | 140 |
| Water |  | - | 112 |
| Other |  | - | 153 |
| $B$-factors (Å<sup>2</sup>) | | | |
| Protein |  | - | 44 |
| Ligand |  | - | 43 |
| Water |  | - | 37 |
| Other |  | - | 55 |
| r.m.s. deviations |  |  |  |
| Bond lengths (Å) |  | - | 0.008 |
| Bond angles (°) |  | - | 0.87 |

<sup>1</sup> Anisotropic resolution limits along each of the three principal directions as defined by AIMLESS based on $Mn(I/sd(I)) > 2$ . Values in parentheses are resolution limits in each direction based on half dataset correlation $> 0.5$ ( $CC_{1/2}$ ).
<sup>2</sup> Nominal resolution is defined based on overall $Mn(I/sd(I)) > 2$ as estimated by AIMLESS.
<sup>3</sup> Values in parentheses are for highest resolution shell

Overall ELOVL7 has seven transmembrane (TM) helices (TM1-TM7) with TM2-7 forming a six TM inverted barrel surrounding a narrow tunnel (Fig. 1b-f), formed by two units of three helices (TM2-4 and TM5-7), each forming an antiparallel three helix arrangement. The two units are assembled as an inverted repeat around the central tunnel (Fig. 1f) with TM1 lying against TM3/4, outside the barrel. This fold is unlike the GPCR 7TM fold and we did not find any six or seven TM protein structures with a similar fold (DALI^20^) (Extended Data Fig. 2). The 35 Å long central tunnel has a narrow (8-10 Å wide) opening on the cytoplasmic face of the protein and is sealed at its ER lumen end by the short buried loop between TM4-5, which connects the two halves of the barrel (Fig. 1b-e). This loop is separated from the ER lumen by the two disulphide-linked ER loops between TMs2-3 and TMs6-7.

### ELOVL7 copurifies with a bound acyl-CoA

Surprisingly, the electron density maps clearly showed a long hooked density from the closed ER end of the tunnel to the cytoplasmic surface of the protein. This electron density is consistent with there being a bound acyl-CoA within the active site tunnel spanning its entire length (Fig. 2a,b and Extended Data Fig. 3). Denaturing intact mass spectrometry analysis of protein samples taken during purification showed two species: an apo form with the mass of the purified protein construct (34132.78 Da) and a form with a mass adduct of 1073.66 Da (Fig. 2c and Extended Data Fig. 1g,h). This mass adduct was observed both with protein obtained from over-expression in *Spodoptera frugiperda* (*Sf*9) insect cells or mammalian (Expi293F) cells. The form with the mass adduct was more stable during purification, so that following size exclusion chromatography (SEC), it represented >90% of the purified protein (Fig. 2c). Based on the electron density and the observed mass adduct, we have modelled a covalently bound 3-keto eicosanoyl-CoA (Mw 1074.02 Da), an analogue of the reaction product which co-purified with ELOVL7. The electron density clearly shows that the adduct is covalently attached to the imidazole rings of both H150 and H181 (Fig. 2d and Extended Data Fig. 3); the C5 carbon of the elongated 3-keto FA is attached to the Nε of H150 and the C2 carbon atom is attached to the Nε of H181. When we introduced a H150A mutation we were not able to detect any adducts (Fig. 3a,b). In contrast, with a H181A mutation we detected both unmodified protein and adducts of 1032.51 and 1059.29Da., consistent with covalent attachment of the acyl-CoAs C18:0 -CoA or C20:1 -CoA, respectively (Fig. 3c).

**Fig. 2.**
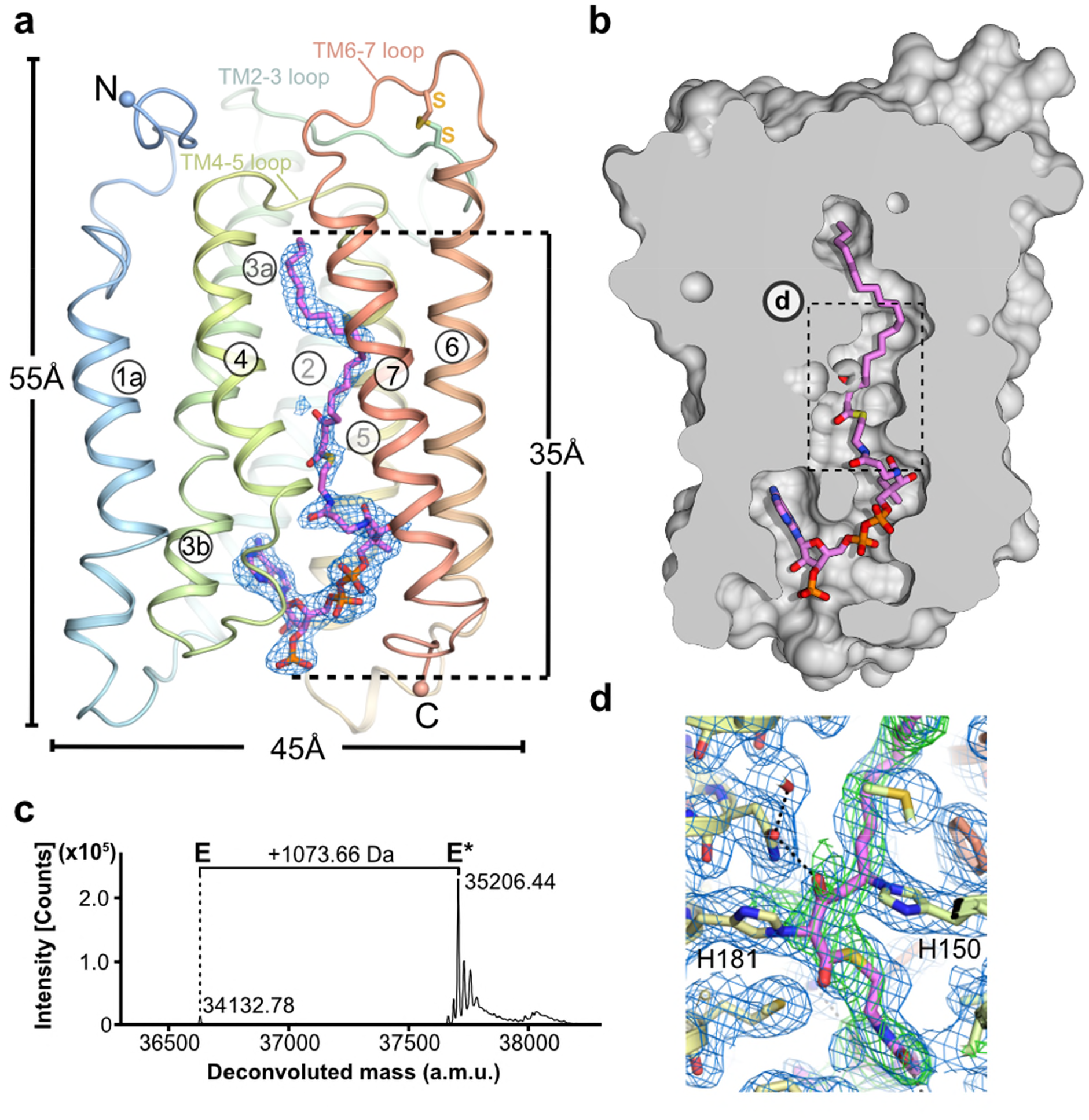
Heterologously expressed ELOVL7 is covalently bound to a 3-keto acyl-CoA. **a**, Cartoon showing the covalently bound 3-keto eicosanoyl CoA along with the FoFc omit electron density map (blue mesh, contoured at 3sigma). **b**, Cutaway molecular surface representation showing the extent of the enclosed central active site tunnel. **c**, Intact mass analysis of purified protein (E) highlighting adduct (E*; +1074Da). **d**, Electron density in region around covalent linkages to active site histidines. Final BUSTER 2mFo-DFc (blue mesh, contoured at 1sigma) and omit mFo-DFc (green mesh, contoured at 2.5sigma) electron density maps are overlaid on the final model.

**Fig. 3.**
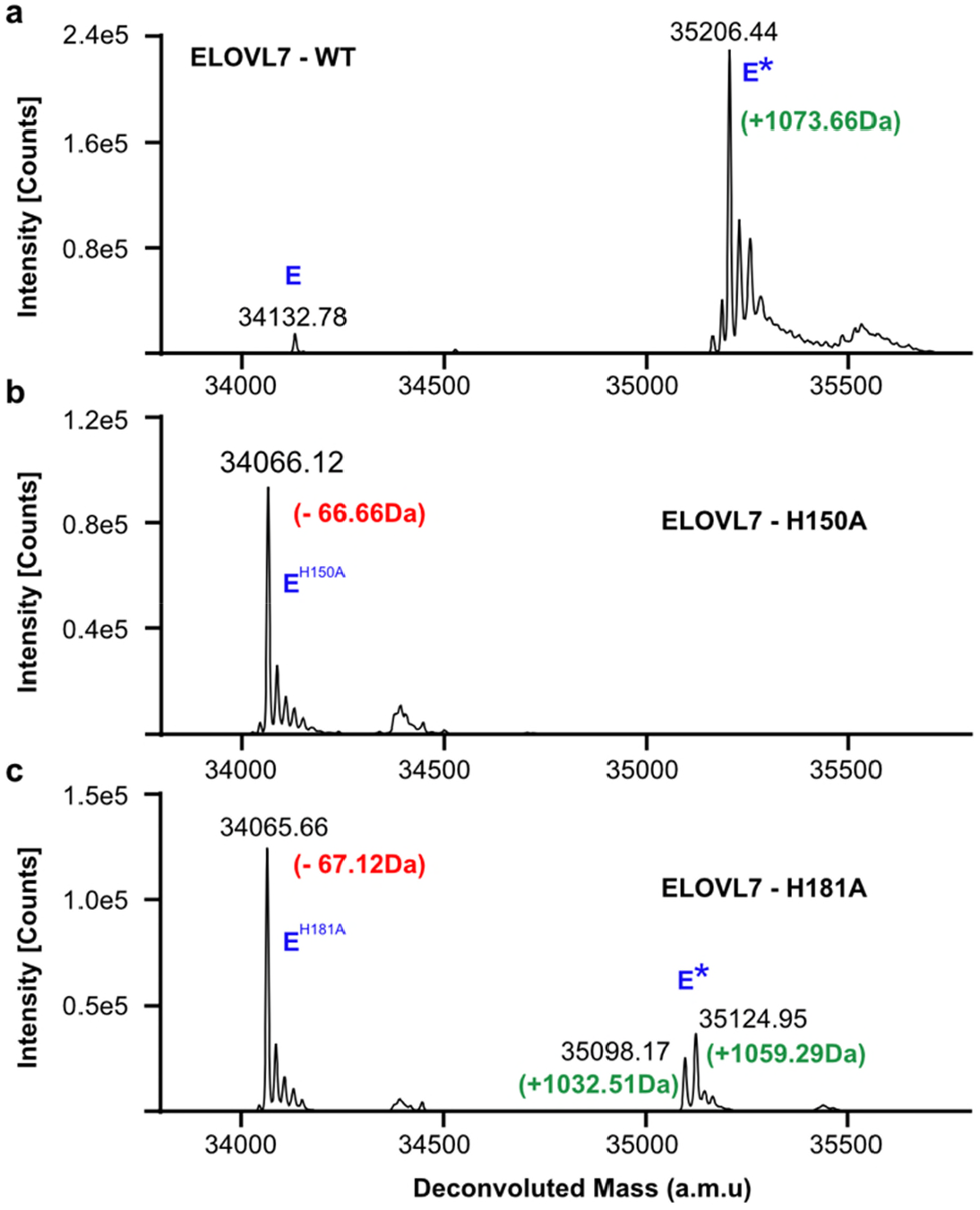
Intact mass analysis of WT ELOVL7, H150A and H181A mutants. **a-c**, Comparison of the deconvoluted intact mass spectra for the untagged **a**, WT enzyme **b**, H150A and **c**, H181A mutant. The expected mass decrease of a His-to-Ala mutation is 66.06 Da. *In vivo* modification (E*) is observed for the WT and H181A mutant but not for H150A.

ELOVLs catalyse the first step of the LCFA and VLCFA elongation cycle through a condensation reaction between an acyl-CoA and malonyl-CoA to yield a 3-keto acyl-CoA^21^. This product then undergoes a series of reduction and dehydration reactions catalysed by three other enzymes to yield an acyl-CoA with two additional carbons in the acyl chain (Fig. 1a). We have confirmed that detergent-purified ELOVL7 converts the substrates stearoyl-CoA (C18:0) and malonyl-CoA to give the product 3-keto-eicosanoyl (C20)-CoA, by mass spectrometry (Extended Data Fig. 4a,b), indicating that although ∼90% of the purified enzyme is covalently modified by the 3-keto acyl-CoA and therefore inhibited, the remaining unmodified protein is active.

### The substrate binding site lies within an extended tunnel

The covalently bound acyl-CoA adduct seen in the crystal structure conveniently marks out the binding sites for both the acyl and CoA components of the substrates and products (Fig. 4), and reveals the architecture of the active site (Fig. 5). The acyl chain binding site is at the upper, occluded end of the tunnel and is terminated by very short TM4-5 loop (Fig. 4a,b). It is lined predominantly by hydrophobic sidechains (W158, G161, G170, F238 and I242), and is curved, allowing both unsaturated and saturated acyl chains to bind (Fig. 4b,c). The acyl chain of the 3-keto acyl-CoA seen in the structure has 20 ordered carbons and it does not occupy the entire pocket, with space for 2 additional carbons, suggesting that 3-keto acyl-CoA products with up to 22 carbons could be accommodated. ELOVL7 has a preference for C18-CoA substrates; it can accept fatty acyl-CoA substrates with carbon chains up to C20 in length ^3,21^, consistent with the observed acyl chain length in the structure. The seven ELOVLs have different preferences for acyl chain length and number of double bonds ^3^, and sequence alignments of the human ELOVL elongases suggest that the residues lining the acyl chain binding site are not well conserved (Fig. 4c; Extended Data Fig. 5b), reflecting their abilities to accommodate different acyl chains.

**Fig. 4.**
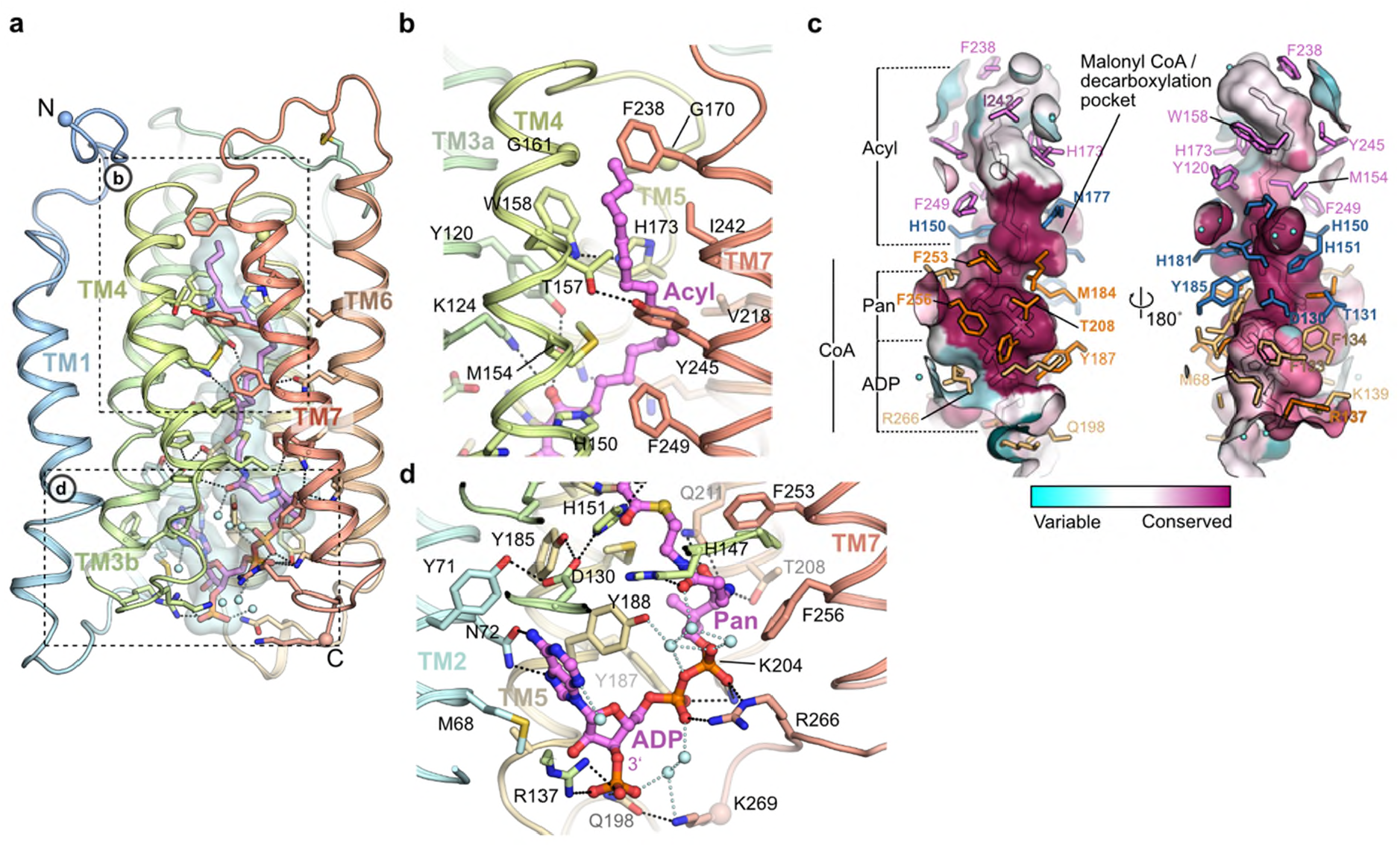
Acyl chain and CoA binding sites. **a**, Overview of the central acyl-CoA binding tunnel. **b**, Acyl chain binding pocket. **c**, Conservation of active site tunnel. Molecular surface representation is coloured by amino acid conservation score calculated by CONSURF^49^ analysis of a diverse set of ELOV1-7 family members. The various subregions of the tunnel are indicated (ADP / Pan from CoA and Acyl chain). Amino acid residues that form the binding tunnel are coloured according to region (pink, acyl; blue, catalytic site; orange CoA binding). **d**, Cytoplasmic-facing CoA 3’-phospho ADP/pantothenic (Pan) binding pocket.

**Fig. 5.**
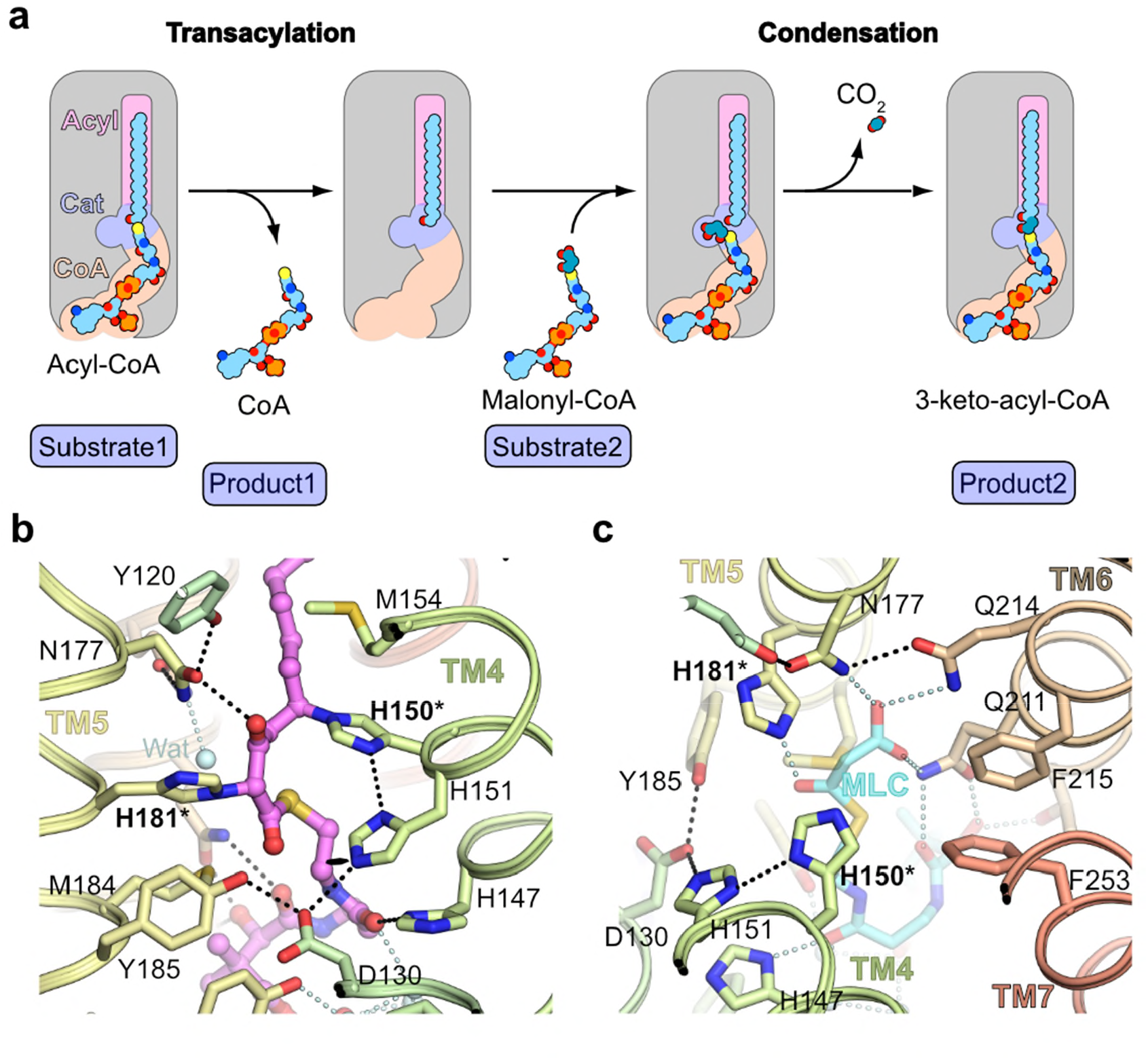
Active site. **a**, Schematic outlining proposed ELOVL ping-pong mechanism. **b-c**, Detailed view of **b**, central catalytic site around conserved histidine motif and **c**, putative decarboxylation pocket. Covalently modified histidines (H150/H181) are highlighted by asterisks. Bound malonyl-CoA (cyan; MLC) has been modelled based on the coordinates of the acyl-CoA.

The CoA binding site lies at the open, cytoplasmic end in the tunnel, with the CoA almost completely buried within the core of the enzyme (95% of the 1400 Å^2^ ligand surface area is buried). The 3’-phospho-ADP/pantothenate portion of CoA adopts a ‘U’ shaped conformation wrapped around the sidechain of Y188 (Fig. 4d). The adenine ring is inserted in a narrow cleft between TM2, 3 and 5. N72 on TM2 forms hydrogen bonds to both N5 and N6 on the adenine base and D130 on TM3b provides a third hydrogen bond to the base (Fig. 4d). The 3’ phosphate projects towards the solvent and forms a salt bridge with R137. The diphosphate forms salt bridges with K204 and R266 and a hydrogen bond with the sidechain of Y187. The pantothenate (Pan) moiety follows a helical path and is hydrogen bonded to the sidechains of T208, Q211 and H147. The cysteamine is elongated and sits in a hydrophobic pocket surrounded by M184 and F253. Together these extensive interactions ensure a highly specific binding site for the CoA portion of the substrates.

### Fatty acid extension proceeds via a ping-pong type mechanism

Acyl transferase reactions generally involve either a ping-pong or a ternary complex mechanism. In the first, the two substrates bind one after the other and a covalent adduct is involved, whereas in the second both substrates bind at the same time and the acyl group is transferred directly from one substrate to the other. The narrow substrate binding tunnel does not provide sufficient space for both substrates to bind at the same time (Fig. 2b), excluding a ternary complex mechanism. The narrow tunnel is however consistent with a ping-pong type mechanism, where the first product must leave before the second substrate binds (Fig. 5a). Therefore we propose that in the first step the acyl-CoA binds, with the FA tail occluded at the closed ER end of the tunnel and the CoA moiety near the entrance. Then the CoA is cleaved and can dissociate, leaving behind the buried acyl chain. In the second step the malonyl-CoA binds at the more accessible CoA binding site, with the malonyl unit in a side pocket in the central active site. Decarboxylation and carbon-carbon bond formation will then yield the 3-keto acyl-CoA product. The entry and exit route for acyl-CoA and 3-keto acyl-CoA is likely to involve lateral movement of the hydrophobic acyl chain directly into the membrane, between two TM helices, rather than having the acyl chain move along the whole of the tunnel, parts of which are hydrophilic. The structure suggests that this entry/exit portal lies between TM4 and TM7, between the two 3TM halves of the barrel (Fig. 1f).

The catalytic site of ELOVL elongases lies at the centre of the membrane and the residues lining this section of the tunnel are highly conserved across all seven elongases (Fig. 4c). A site directed mutagenesis study in the ELOVL homologue EloA from the social amoeba *Dictyostelium discoideum* identified a series of residues that were essential for activity (equivalent to K124, D130, H150, H151, N177, H181 and Q211 in ELOVL7) ^22^ and all of these residues lie within the active site. The active site is lined with histidine residues (Fig. 5b,c), including the canonical ELOVL elongase HxxHH motif ^21^. Although this type of histidine motif is sometimes associated with metal ion binding, for example in fatty acyl desaturases ^23^, the electron density maps show no indication of bound metal ions and the geometry is not suitable for metal ion binding, indicating that metal-assisted catalysis is unlikely. Although cysteines are often involved in acyl transferase reactions, there are no cysteine residues in or near the active site that could be involved in catalysis. The structure shows the CoA thiol sits just below a triad of histidines – H150 and H151 on TM4 from the conserved HxxHH motif and H181 on TM5, all conserved residues required for ELOVL activity ^22^ (Fig. 5b). In the crystal structure H150 and H181 are covalently attached to the copurified product analogue, and mutation of these residues affects the *in vivo* modification of heterologously expressed ELOVL7. The H150A mutation gave only unmodified protein, suggesting that H150 could be the first residue to be modified, whereas the H181A mutation resulted in modification by substrate analogues, suggesting that H181 could be involved in the subsequent catalytic steps (Fig. 3b,c).

### ELOVL7 catalysis proceeds via an acyl-imidazole intermediate involving H150

Using the ELOVL7 structure with the bound product analogue we modeled the positions of each substrate and product during the reaction, indicating how the reaction could proceed (Fig. 6a-c). Once the first substrate, the fatty acyl-CoA has bound, then H150 is ideally placed to act as a nucleophile to attack the carbonyl group of the fatty acyl-CoA. The nucleophilicity of H150 is enhanced by a bonding network involving H151 and D130 that ensures that the Nε atom is unprotonated (Fig. 6a). The sidechains of N177 and H181, together with a water molecule (which hydrogen bonds to N177 and Q214), are suitably positioned to stabilise the oxyanion prior to release of CoA. This transient tetrahedral intermediate would then collapse back to the carbonyl, ejecting the CoA, which could acquire a hydrogen atom from H181, providing protonation of the CoA thiol and completing the transacylation step of the reaction (Fig. 6b). The formation of a covalent acyl-enzyme intermediate at the end of the first step was confirmed by denaturing intact mass spectrometry analysis of the enzyme after incubation with an acyl-CoA substrate (C18:0-CoA). We observed a 266.51 Da adduct (Fig. 6d and Extended Data Fig. 6a,b), consistent with the transfer of a C18:0-acyl group to the protein. However, incubation with both substrates, C18:0-CoA and malonyl-CoA, greatly reduced accumulaton of the adduct intermediate, consistent with the reaction having gone to completion (Fig. 6d and Extended Data Fig. 6c). Similarly, incubation with the unsaturated C18:3(n3)-CoA substrate alone resulted in the formation of a 260.74 Da adduct, whereas incubation with C18:3(n3)-CoA and malonyl-CoA did not give an adduct, consistent with covalent addition of the unsaturated acyl chain of the substrate as the first step, followed by completion of the reaction with the second substrate (Extended Data Fig. 6f,g). Addition of EDTA or EGTA did not interfere with covalent modification of the protein by either substrate, supporting the hypothesis that metal ions are not required for catalysis (Extended Data Fig. 6d-g).

**Fig. 6.**
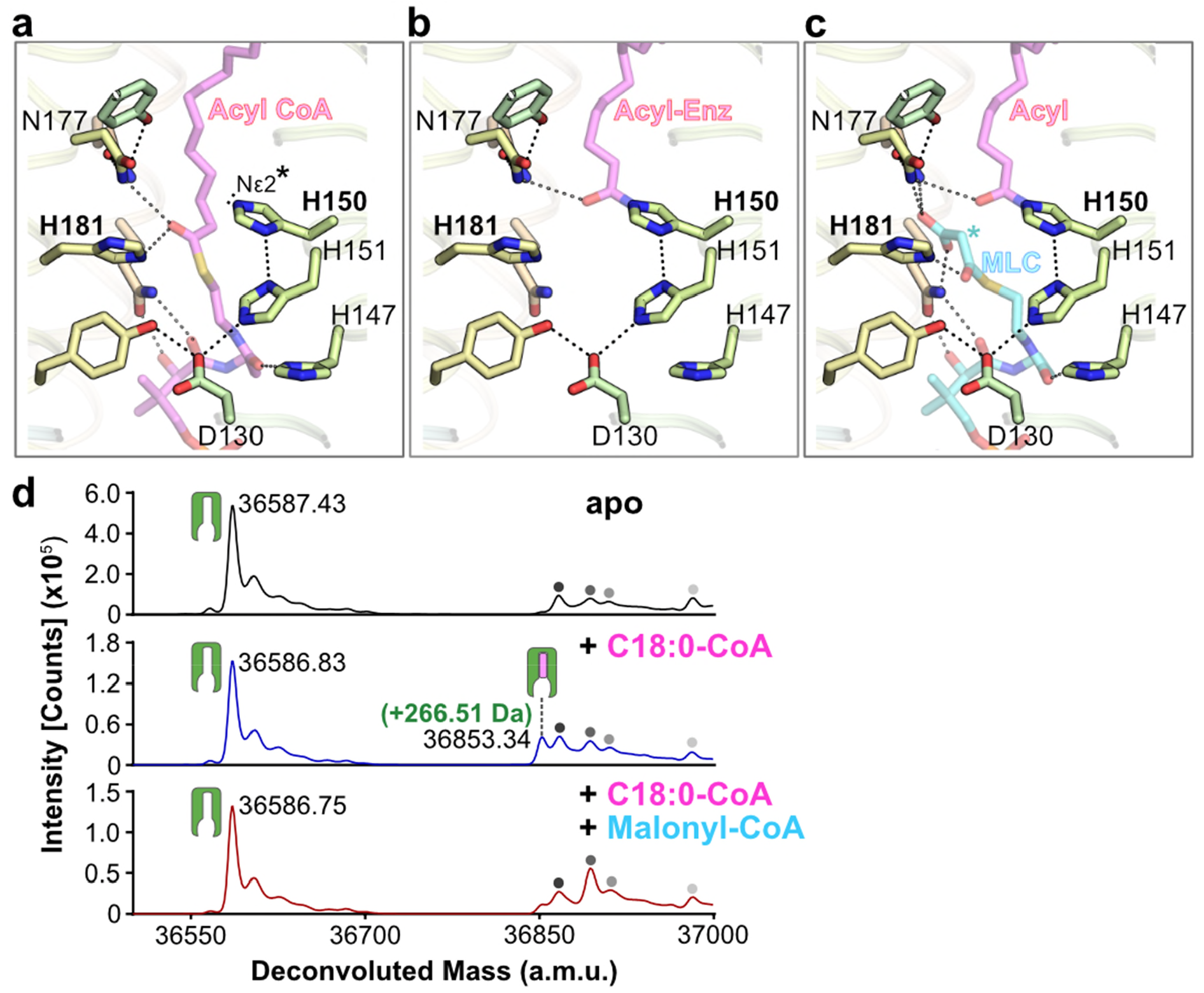
Proposed steps in catalytic mechanism and evidence for a covalent acyl-imidazole intermediate. **a-c**, Proposed catalytic steps for **a**, acyl-CoA binding; **b**, transacylation to form an acyl-imidazole intermediate. **c**, Proposed malonyl-CoA (MLC) condensation step with acyl intermediate. Ligand complexes are models based on product/inhibitor crystal structure. The unprotonated imidazole nitrogen of H150 involved in the transacylation step and the location of the enolate carbon nucleophile formed after decarboxylation of MLC are highlighted with asterisks. **d**, Intact mass analysis of protein only (black), protein + C18:0 acyl-CoA (blue) and protein + C18:0 acyl-CoA + malonyl-CoA (red). Peaks are indicated as follows: protein (green icon), acyl enzyme intermediate (green icon+pink oblong), background species in all traces (grey circles).

Many fatty acid synthases and other acyltransferases require the formation of acyl-enzyme intermediates with either oxy- or thio-ester covalent linkages ^24^, usually involving cysteine or serine ^25^ residues. However, there are no cysteines or serines in or near the ELOVL active site. The structure indicated that H150 was a candidate for this role and a H150A mutation in ELOVL7 gave protein that was not modified upon incubation with C18:3(n3)-CoA (Extended Data Fig. 7a,b) (we were unable to use C18:0-CoA as substrate due to the presence of background peaks in the MS traces). In contrast, incubation of the H181A protein with C18:3(n3)-CoA gave a covalent adduct of 261.97 Da (Extended Data Fig. 7c,d). These results are consistent with the H150 sidechain being the site where the fatty acyl unit is covalently attached. Covalent acyl adducts to histidines are uncommon, but they are not entirely without precedent: HlyC, the enzyme that activates the *Escherichia coli* toxin prohemolysin, catalyses acyl transfer via a histidine ^24,26^ and the enzymes of the complement system, C3 and C4, utilise a histidine for internal acyl transfer ^27,28^. In addition artificial enzyme systems have been created in which a nucleophilic histidine is acylated to form a covalent acyl-imidazole intermediate, confirming that such additions are possible ^29-31^.

In the second ‘condensation’ step, malonyl-CoA is combined with the bound acyl chain through a Claisen condensation. This reaction requires malonyl-CoA binding, decarboxylation to form an enolate, followed by reaction with the acyl-enzyme intermediate to form the 3-keto acyl-CoA product that is elongated by 2 carbon units. Initially malonyl-CoA would bind to the CoA binding site, and modelling suggests that the malonyl unit would be positioned so that the carboxylate would lie in a pocket at the side of the tunnel between TM5 and TM6 (Fig. 5c). This shallow pocket is lined by a semicircle of amide groups from N177, Q211 and Q214, which would hydrogen bond to the carboxylate and, together with H181, could catalyse decarboxylation (Fig. 6c and Extended Data Fig. 8). Modelling suggests that His181 lies adjacent to the thioester oxo group of malonyl-CoA, where it could promote decarboxylation by stabilizing enolate formation. Mutation of these residues in EloA gave either inactive enzyme (H181, N177 and Q211 (ELOVL7 numbering)), or enzyme with 10% of WT activity (Q214). The resulting nucleophilic enolate carbanion would then be well-placed to react with the acyl-CoA adduct on H150, to form the 3-keto acyl-CoA product (Extended Data Fig. 8) which would then be released from the enzyme.

Interestingly, the crosslinked version of the product analogue seen in the crystal structure is covalently linked to the C5 of the 3-keto acyl product analogue inhibitor, not directly the C3-carbonyl. We speculate that this crosslink could have been the result of ELOVL7’s conjugate addition to an unsaturated acyl-CoA substrate or breakdown product, with an adjacent double bond in the acyl chain (e.g. 2,3-*trans*-enoyl-CoA), thus allowing the formation of a displaced acyl-imidazole covalent bond. Acyl-CoAs with a double bond in this location are formed both during the FA elongation cycle and during breakdown of VLCFAs in the peroxisomes.

## Discussion

The structure of ELOVL7 reveals a narrow extended tunnel which is involved in binding and extension of acyl-CoAs. Previous studies have highlighted a series of histidine residues that are essential for ELOVL enzyme activity and these are clustered halfway along the membrane-embedded tunnel, at the midpoint of the bilayer, where they are poised for catalysis. Through a combination of mutagenesis and intact protein MS analysis, our results show that the elongation reaction proceeds through a stable acyl-imidazole intermediate formed between the substrate acyl-CoA and the second His in the ELOVL HxxHH motif. We propose that a second group of sidechains previously demonstrated to be required for elongase activity (N177, H181, Q211, and Q214)^22^ lie adjacent to the modified histidine, where they could play a role in the decarboxylation of malonyl-CoA, the second substrate. The resulting reactive enolate nucleophile could then react with the acyl-enzyme intermediate giving rise to the 3-keto acyl-CoA product.

Failure of the VLCFA elongation process through ELOVL mutations leads to loss of ceramides and sphingolipids required for myelin, skin barrier and retinal function. Mutations in the ELOVL4 gene cause Stargardt disease-3^10^, ichthyosis, intellectual disability, spastic quadriplegia^32^, and spinocerebellar ataxia 34 (SCA34)^11,33^. ELOVL5 mutations could cause spinocerebellar ataxia 38 (SCA38)^12^. *Elovl3* knockout mice suffer severely from hair loss and have an imbalance in the lipid species of the sebum^34,35^; and mouse knockout studies of ELOVL5 / ELOVL6 suggest associations with hepatic steatosis^13^ and obesity-induced insulin resistance ^14^. ELOVL7, the most recently discovered ELOVL elongase, is associated with prostate^15,16^ and gynaecological ^17^ cancer and early onset Parkinson’s disease ^18^. ELOVL7 knockdown reduces cell death and membrane permeabilization during necroptosis, a form of programmed cell death^19^. The unusual architecture, active site and chemistry revealed by our ELOVL elongase structure provides exciting new avenues for design of modulators of ELOVLs. These may be of value for patients with X-linked adrenoleukodystrophy (X-ALD), as they are unable to break down VLCFAs, leading to accumulation in cells and further elongation by ELOVL1^36^. X-ALD affects the adrenal cortex and nervous system, leading to adrenocortical insufficiency and myelopathy, with progressive demyelination of nerves in severe cases. The formulation known as Lorenzo’s oil has been shown to alleviate symptoms of X-ALD and is reported to inhibit ELOVL1^37^. The design of specific ELOVL elongase inhibitors may offer a route to reduction of VLCFAs, thus providing a novel approach to therapy for this devastating disease.

## Methods

### Cloning

The *Homo sapiens ELOVL7* gene, encoding the ELOVL7 protein (Uniprot ID: A1L3×0) was cloned into the baculovirus transfer vector pFB-CT10HF-LIC (available from The Addgene Nonprofit Plasmid Repository) for expression in *Sf9* cells (Thermo-Fisher Scientific, 11496015) using the primers shown in Extended Data Table 1. A C-terminal TEV-cleavable His10-FLAG tag was fused to the protein for purification. For mammalian expression, the same construct was cloned into the pHTBV1.1-LIC baculovirus transfer vector (The BacMam vector backbone (pHTBV1.1), kindly provided by Professor Frederick Boyce, Massachusetts General Hospital, Cambridge, MA and adapted for ligation independent cloning in-house for expression in Expi293F cells (Thermo-Fisher Scientific) which similarly confers a TEV cleavable C-terminal His10-FLAG tag.

The mutagenesis was performed using Q5^®^ Site-Directed Mutagenesis Kit (NEB) according to manufacturer’s instructions using primers listed in Extended Data Table 1.

### Protein Production

For both insect and mammalian expression, baculoviral DNA was produced by transposition of DH10Bac with either the ELOVL7-pFB-CT10HF-LIC or ELOVL7-pHTBV1.1-LIC transfer vectors, which were then used to transfect *Sf9* cells to produce baculovirus for transduction using the transfection reagent Insect GeneJuice® (MerckMillipore). The virus amplification was performed by infecting mid-log *Sf9* cells (2 × 10^6^ cells ml^-1^) grown in *Sf*-900II™ media with 2% fetal bovine serum. For large scale protein production, *Sf9* cells at the density of 2 × 10^6^ cell ml^-1^ were infected with 5 ml of P2 (second passage) recombinant baculovirus in *Sf*-900™ II in 2 L roller bottles (Biofil) and incubated for 72 h at 27°C. Cells were harvested by centrifugation at 900 x *g* for 15 mins, washed with phosphate buffered saline (PBS), and pelleted again prior to flash freezing in liquid N_2_, then stored at −80 °C for further use.

For mammalian (Expi293F) expression, 1 L of Expi293F cell cultures (2 × 10^6^ cells ml^-1^) in Freestyle 293™ Expression Medium (Thermo-Fisher) were transduced with 30 ml of P3 baculovirus (third passage) in the presence of 5 mM sodium butyrate in a 2 L roller bottle (Biofil). Cells were grown in a humidity controlled orbital shaker for 48 hours at 37 °C with 8% CO_2_ before being harvested using the same process as for insect cells.

### Protein Purification

All the following steps were performed at 4°C unless otherwise indicated. Cell pellets were resuspended in lysis buffer (50 mM HEPES-NaOH, pH 7.5, 500 mM NaCl, 5% v/v glycerol, 1 mM TCEP-NaOH, Roche protease inhibitor cocktail EDTA-free) at the ratio of 50 ml / L equiv. original cell culture. The resuspension was then passed twice through an EmulsiFlex-C5 homogenizer (Avestin) at 10000 psi. Membrane proteins were extracted from the cell lysate with 1% w/v octyl glucose neopentyl glycol (OGNG; Generon, Cat. No. NG311) / 0.1% cholesteryl hemisuccinate tris salt (CHS; Sigma-Aldrich, Cat. No. C6512) and rotated for 2 h. Cell debris was removed by centrifugation at 35,000 x *g* for 1 h. The supernatant was supplemented with 5 mM imidazole pH 8.0 before incubation with Co^2+^ charged TALON resin (Clontech) for 1 h on a rotator (1 ml resin slurry per L original culture volume). The Talon resin was collected by centrifugation at 700 x *g* for 5 mins and washed with 30 column volumes of washing buffer (50 mM HEPES-NaOH, pH 7.5, 500 mM NaCl, 1 mM TCEP-NaOH, 0.12% w/v OGNG / 0.012% w/v CHS and 20 mM imidazole pH 8.0) before the target protein was eluted with elution buffer (washing buffer supplemented with 250 mM imidazole pH 8.0). The eluted protein was then exchanged into lysis buffer supplemented with 0.15% w/v OGNG / 0.015% w/v CHS by passing over a pre-equilibrated Sephadex PD-10 desalting column (GE Life Sciences). TEV protease was added to the desalted protein at a weight ratio of 1:10 for overnight tag cleavage. The His-tagged TEV protease was removed by a second Talon resin binding step for 1 h, and the flow through was collected and concentrated to <1ml using a 100 kDa molecular weight cutoff (MWCO) Vivaspin 20 centrifugal concentrator (GE Life Sciences) and further purified by size exclusion chromatography on a Superdex 200 10/300 Increase GL column (GE Healthcare) in size exclusion chromatography (SEC) buffer (20 mM HEPES-NaOH, pH 7.5, 200 mM NaCl, 1 mM TCEP-NaOH, 0.08% w/v OGNG/ 0.008% w/v CHS). The H150A and H181A mutated protein were expressed and purified in the same way as the wild type protein. In addition we attempted to produce a His151Ala version of the protein, but this variant proved to be too unstable to purify.

For structural studies, the flow through from the reverse Talon step was incubated with 50 mM iodoacetamide (IAM) (Merck Millipore) for 20 mins at room temperature. IAM was removed by passing the reaction mixture down a PD-10 desalting column prior to concentration. After SEC, fractions containing the highest concentration of ELOVL7 were pooled and concentrated to 12-25 mg/ml using a 100 kDa MWCO Vivaspin 20 centrifugal concentrator.

### Crystallization

Initial protein crystals were grown at 4°C in condition E10 of the MemGold2-ECO Screen (Molecular Dimensions; 0.05 M Na-acetate pH 4.5, 0.23 M NaCl, 33 % v/v polyethylene glycol (PEG) 400) in 3-well sitting-drop crystallisation plates (SwissCi) with 150 nl drops and 2:1 and 1:1 protein to reservoir ratios. Crystals appeared after 4-7 days and grew to full size within 3-4 weeks. Two microlitre hanging drops were set up in 24-well XRL plates (Molecular Dimensions) at protein to reservoir ratios of 2.5:1, 2:1 and 1.5:1. The best crystals grew in 0.1 M Na-acetate pH 4.5, 0.23 M NaCl, 34-38% v/v PEG400 using the IAM-modified protein at a concentration of 5-8 mg/ml and were harvested after 12-14 days. Crystals could also be grown for the unmodified native protein but were limited in size and poorly reproducible despite using seeding. Prior to vitrification, crystals were sequentially transferred to mother liquor solutions with an increasing amount of PEG400 to a final concentration of PEG400 of 46% v/v over 10-15 mins. For heavy atom derivatisation, crystals were looped into drops containing reservoir solution supplemented with 10 mM mercury chloride and soaked for 10 mins. Hg-soaked crystals were then treated to the same PEG400 escalation strategy using Hg-free solutions before being vitrified in liquid nitrogen.

### Data Collection

All crystals were screened and X-ray diffraction data were collected on the I24 microfocus beamline at the Diamond Light Source (Didcot, UK) from vitrified crystals at 100 K. Multiple data sweeps were collected from crystals using 0.2° oscillation and a beam-size of 20 μm x 20 μm. Crystals were translated by at least 25 μm between collection areas. Native data were collected at a wavelength of 0.9686 Å. The high resolution native dataset was assembled from four overlapping 100° wedges of data collected from separate volumes of a single crystal. 960° of anomalous data were also collected from a mercury-derivatised crystal at a wavelength close to the Hg L_III_ edge (λ = 0.992 Å) in a single pass.

Unmerged scaled data from the *xia2*.*dials* automated processing pipeline^38^ was truncated and merged using AIMLESS (CCP4^39,40^). ELOVL7 crystals are monoclinic and contain two copies of the enzyme in each asymmetric unit. All diffraction data were highly anisotropic and limited to between 3.4 - 4.5 Å resolution in the worse direction and 2.05 – 3 Å in the best direction (Table 1). Data truncated in AIMLESS with isotropic resolution limits was used for phasing and initial cycles of model building. The high resolution native dataset (nominal overall resolution 2.6 Å) was further processed with STARANISO (v1.0.4^41^) to the maximum resolution of 2.05 Å and used for refinement (see Table 1 for details).

### Phasing

Phasing was carried out in PHENIX^42^ using SIRAS with the Hg-peak data and a 3Å isomorphous lower resolution native dataset. Two Hg^2+^ sites were located with *phenix*.*hyss* using data to 4.5 Å. The resulting 3 Å phased electron density map had clear protein density allowing the identification of the NCS relationship between the two ELOVL7 molecules in the asymmetric unit. After two-fold averaging using RESOLVE^43^, the resultant map was of sufficient quality to manually model all the TM helices. Initial phases were further improved by cross-crystal averaging with a non-isomorphous, less anisotropic and slightly higher resolution dataset (unit cell dimensions 64.14 Å x 71.93 Å x 111.74 Å β = 106.7°) using DMMULTI^44^. The resultant map was of excellent quality and the majority of the structure could be built automatically with BUCCANEER^45^.

### Model Building and Refinement

The BUCCANEER-built model was manually rebuilt in COOT^46^ and refined using BUSTER (version 2.10.3, 29-NOV-2019^47^). The model was refined with LSSR restraints^48^ and a single TLS group was refined for each protein chain. Ligand dictionaries were generated using GRADE (Global Phasing Ltd). All data to 2.05 Å from the STARANISO processing was used in refinement despite the low completeness and truncation to lower resolutions resulted in a refined model with worse geometry. The final model comprises residues 16-269 (chain B, 14-269), a covalently bound 3-keto-CoA acyl lipid, four OGNG detergent molecules and 112 solvent molecules. None of the resolved cysteines appear to be modified with IAM. Data collection and refinement statistics are reported in Table 1. Molecular models for the complexes with substrate, covalently bound acyl-imidazole intermediate, malonyl-CoA and 3-keto-eicosanoyl product were built in COOT based on the crystal structure using simple manual placement and correction of any steric clashes.

### Denaturing Intact Mass Spectrometry

The denaturing intact mass spectrometry measurements were performed using an Agilent 1290 Infinity LC System in-line with an Agilent 6530 Accurate-Mass Q-TOF LC/MS (Agilent Technologies Inc.). The solvent system was consisted of 0.1% Optima™ LC/MS grade formic acid (Fisher Chemical) in HPLC electrochemical grade water (Fisher Chemical) (solvent A) and 0.1% formic acid in Optima™ LC/MS grade methanol (Fisher Chemical) (solvent B). Typically, 1-2 μg of protein sample was diluted to 60 μl with 30% methanol in 0.1% formic acid. 60 μl of sample was injected onto a ZORBAX StableBond 300 C3 column (Agilent Technologies Inc.) by an auto sampler. The flowrate of the LC system was set to 0.5 ml/min. 30% of solvent B was applied in the beginning and the sample elution was initiated by a linear gradient from 30% to 95% of solvent B over 7 min. 95% B was then applied for 2 min, followed by 2 min equilibration with 30% B. The mass spectrometer was in positive ion, 2 GHz detector mode and spectra were recorded with capillary, fragmentor and collision cell voltages of 4000 V, 250 V and 0 V, respectively. The drying gas was supplied at 350°C with flow rate of 12 l/min and nebulizer at 60 psi. The data was acquired from 100-3200 m/z. Data analysis was performed using MassHunter Qualitative Analysis Version B.07.00 (Agilent) software.

Assignment of the deconvoluted mass peak to the apo form of the protein was achieved by identifying a −89.62 Da mass shift relative to the theoretical mass of the purified protein. This corresponds to the loss of the initiator methionine, followed by acetylation of the new N-terminus (theoretical mass shift of −89.16 Da). Further characterisation of adduct formation and IAM modification was carried out by analysing the deconvoluted mass shift from the apo protein peak to those corresponding to covalently modified forms of the protein. This enabled the determination of a mass shift of +1073.66 Da for the copurified adduct, and the observation that 1, 2 or 3 sites can be simultaneously modified by IAM (predicted +57.07 Da per site modified).

In order to trap the covalent acyl-enzyme intermediate, the purified, tagged, wild-type ELOVL7 protein at 1.5 mg/mL (obtained after the desalting step that followed IMAC elution) was incubated with 100 μM C18:0-CoA (Avanti Polar Lipids, Cat. No. 870718) or C18:3(n3)-CoA (Avanti Polar Lipids, Cat. No. 870732) for 2 hours at 37 °C, in the presence and absence of 1 mM EDTA, 1 mM EGTA, or 100 μM Malonyl CoA (Sigma-Aldrich, Cat. No. M4263). The reaction was terminated by dilution into 30% methanol in 0.1% formic acid, as described above. Covalent acyl-enzyme intermediate formation was identified by monitoring the presence of a mass shift upon incubation with the substrate, corresponding to the addition of the substrate acyl chain through attachment of the histidine imidazole to the thioester carbonyl, resulting in thioester cleavage and loss of CoA (predicted +266.47 Da upon reaction with C18:0-CoA; +260.42 Da upon reaction with C18:3(n3)-CoA). The site of covalent modification was probed by carrying out this experiment with the His150Ala and His181Ala mutants, which allowed identification of His150 as the nucleophile involved in covalent intermediate formation.

### Product formation detected by mass spectrometry

To follow the enzymatic reaction, a small molecule mass spectrometry experiment was performed using LC-MS with a Waters system equipped with a Waters 2545 binary gradient module, a Waters SQ Detector 2, Waters 2489 UV/visible detector, and a Waters 2424 ELS detector. Masslynx 4.0 software by Waters (Beverly, MA) was applied for data processing. Analytical separation of malonyl-CoA, C18:0-CoA and the 3-keto-eicosanoyl (C20:0)-CoA reaction product was carried out on a Phenomenex Kinetex 5 μm EVO C18 column (100 mm × 3.0 mm, 100 Å) using a flow rate of 2 mL/min in a 3 min gradient elution, with 20 μL injections. The mobile phase was a mixture of 93% H_2_O, 5% acetonitrile, and 2% of 0.5 M ammonium acetate adjusted to pH 6 with glacial acetic acid (solvent A) and 18% H_2_O, 80% acetonitrile, and 2% of 0.5 M ammonium acetate adjusted to pH 6 with glacial acetic acid (solvent B). Gradient elution was as follows: 95:5 (A/B) 0.35 min, 95:5 (A/B) to 5:95 (A/B) over 1 min, 5:95 (A/B) over 0.75 min, and then reversion back to 95:5 (A/B) over 0.1 min and 95:5 (A/B) over 0.8 min. A blank injection of MeOH (20 μL) was included between runs to avoid any carry-over from the previous sample.

Samples were analysed by mass spectrometry in negative ESI single ion scans set for mass ions of 852.4 m/z (ESI-) (malonyl-CoA, Sigma-Aldrich, Cat. No. M4263), 1032.9 m/z (ESI-) (C18:0-CoA, Sigma-Aldrich, Cat. No. S0802) and 1074.9 m/z (ESI-) (3-keto-eicosanoyl-CoA). The retention times for these compounds were 0.23, 1.60 and 1.63 mins, respectively. The capillary voltage was set to 3.00 kV, the cone voltage was set to 30 V. Nitrogen was used as desolvation gas at flow rate of 500 L/h. Desolvation temperature was set to 250 °C.

### Size exclusion chromatography with multiangle light scattering (SEC-MALS)

SEC-MALS analysis was performed by injecting 100 μl of purified protein at 1 mg/mL onto a Superdex 200 10/300 Increase GL column (GE Healthcare), pre-equilibrated in 20 mM HEPES pH 7.5, 200 mM NaCl, 0.1% w/v OGNG / 0.01% w/v CHS, using an OMNISEC RESOLVE system (Malvern). Light scattering and refractive index changes were monitored using an integrated OMNISEC REVEAL multi-detector module (Malvern), and the data was analysed using OMNISEC 5.1 software. To enable calculation of the protein molecular weight within the protein-detergent complex, the detergent d*n*/dc was experimentally determined by generating a standard curve of refractive index peak area vs. detergent concentration. The slope of this curve, corresponding to the detergent d*n*/dc, was 0.127 ml/g. The protein d*n*/dc was assumed to be 0.185 ml/g.

## Data Availability

Atomic coordinates and structure factors for the reported crystal structure are deposited in the protein databank (PDB) under accession code 6Y7F.

## Acknowledgements

L.N., A.C.W.P., G.F.R., A.C., V.C., S.R.B., D.S., T.M., L.S., S.M.M.M., N.B.-B., E.P.C. were members of the SGC, a registered charity (number 1097737) that receives funds from AbbVie, Bayer Pharma AG, Boehringer Ingelheim, the Canada Foundation for Innovation, Genome Canada, Janssen, Merck KGaA, Merck & Co., Novartis, the Ontario Ministry of Economic Development and Innovation, Pfizer, São Paulo Research Foundation-FAPESP and Takeda, as well as the Innovative Medicines Initiative Joint Undertaking ULTRA-DD grant 115766 and the Wellcome grant no 106169/Z/14/Z. T.C.P. is supported by a Wellcome PhD studentship. A.Q. is supported by Wellcome grant no 20289/Z16/Z. The authors would like to thank Diamond Light Source for beam time (BAG proposal mx19301), and the I24 beamline staff for assistance with beam time, crystal screening and data collection. We thank Oliver Smart and Clemens Vonrhein at Global Phasing Ltd for assistance with generating covalent ligand restraints. We acknowledge the use of the UCSF Chimera package from the Resource for Biocomputing, Visualisation, and Informatics at the University of California, San Francisco (supported by NIGMS P41-GM103311). We are grateful to the Membrane Protein Laboratory (Wellcome grant ref (20289/Z16/Z)) at the Research Complex at Harwell for access to SEC-MALS equipment and assistance with these experiments.

## Author Contributions

L.N. purified protein, carried out biophysical characterisations, assisted by V.C. and D.S., and obtained crystals that diffracted to beyond 4 Å resolution. L.N. and A.C.W.P. collected X-ray diffraction data and solved and built the structure. T.C.P. purified protein samples, performed mass spectrometry adduct analysis and SEC-MALS experiments. A.Q. provided access to equipment and assisted in SEC-MALS. G.F.R. assisted with design and execution of the rapid fire mass spectrometry activity assay, supervised by P.E.B.. T.M. assisted with mass spectrometry methods optimisation. S.R.B. and A.C. were involved in the early stages of the project, including design of constructs, optimisation of the protein purification, production and screening of initial crystals and initial MS studies. J.D.L. assisted with early stages of the project, including testing constructs and providing materials. Constructs were screened for expression by L.S. and large scale insect cell expressions were produced by S.M.M.M., supervised by N.B.B. Data were analysed and the paper was written by L.N., T.C.P., A.C.W.P. G.F.R., P.E.B. and E.P.C.. E.P.C. supervised all aspects of the project.

## Author Information

Reprints and permissions information are available at www.nature.com/reprints.

The other authors declare no competing interests.

## Extended Data Figures

**Extended Data Fig.1.**
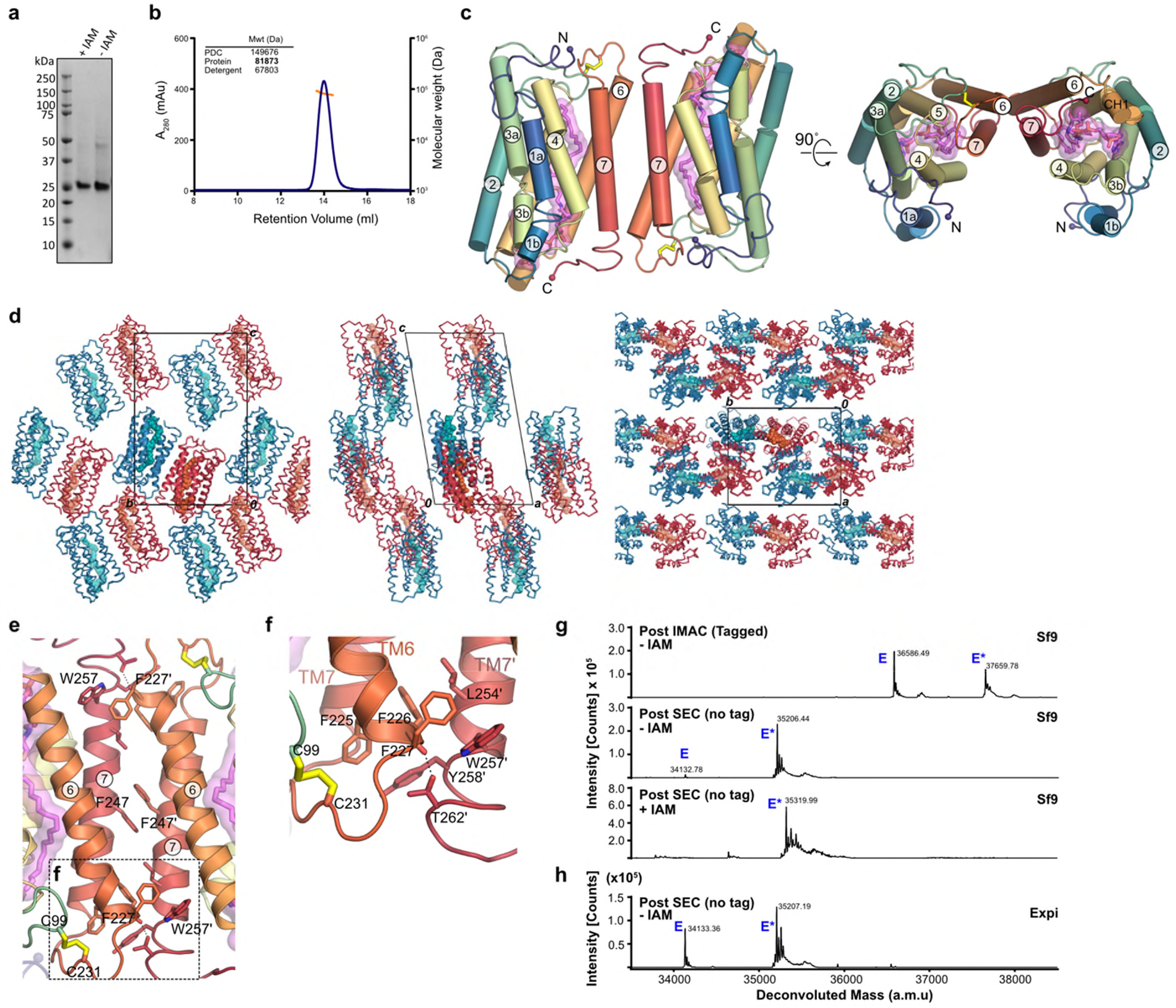
Properties of purified ELOVL7. **a**, SDS-PAGE gel of purified ELOVL7. **b**, SEC profile and MALLS analysis showing that OGNG-solubilised protein exixts as a dimer in solution. **c**, Representation of head-to-tail dimer present in the crystal. **d**, ELOVL7 head-to-tail dimer packing within the crystal lattice. **e, f** Details of dimer interface interactions between TM6 and TM7 of each molecule. **g-h**, Intact mass analysis of ELOVL7 protein at various stages during purification. Deconvoluted mass spectra are shown for ELOVL7 protein purified from (**g**) insect and (**h**) Expi293F cells. For protein purifed after expression in insect cells, the samples are shown after immobilized metal affinity chromatography (IMAC), after cleavage of the tag and size exclusion chromatography (SEC) and after treatment with iodoacetamine (IAM). The expected mass of the untagged ‘apo’ enzyme (E) based on the sequence is 34222 Da. The observed mass peak (34133 Da; −89 Da) corresponds to the loss of the N-terminal methionine (−131.21 Da) and acetylation of the resulting new N-terminus (+42 Da). All samples were run in their reduced state. The modified material (E*) appears as an adduct with an average mass shift of +1073.6 Da. The addition of 113.55 Da upon treatment with IAM suggests modification of two cysteine residues.

**Extended Data Fig.2.**
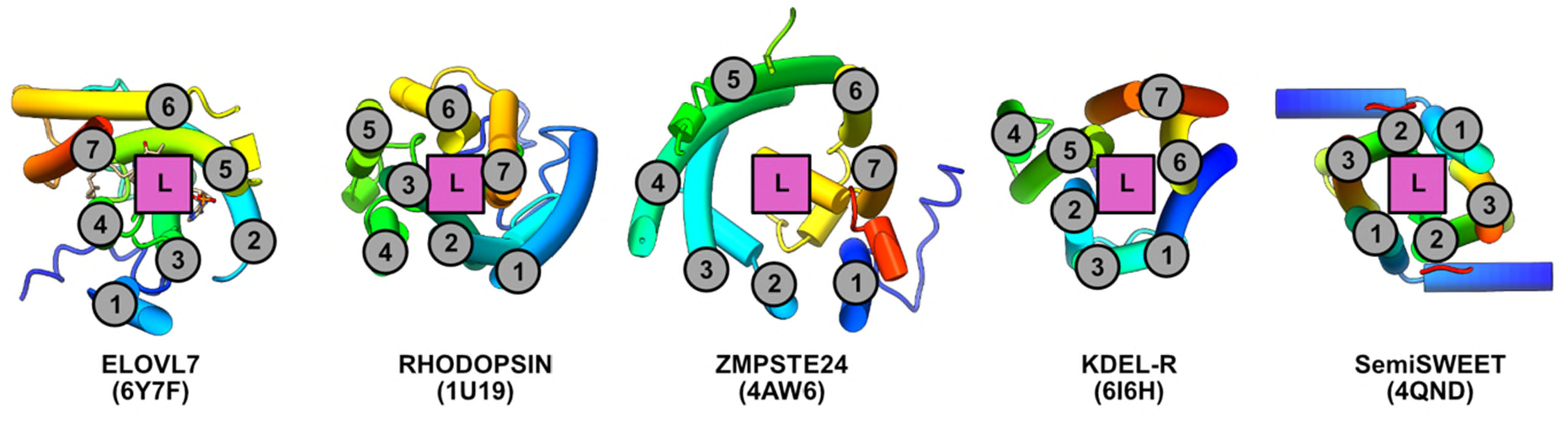
TM helix topology of ELOVL7. TM helical topology of ELOVL7 is compared with other six and seven membered TM bundles. TM helices are numbered and location of substrate/ligand site marked. Underlying cartoon representations of each structure are coloured from blue to red from the N- to C-termini respectively. PDB accession codes are shown in parentheses.

**Extended Data Fig.3.**
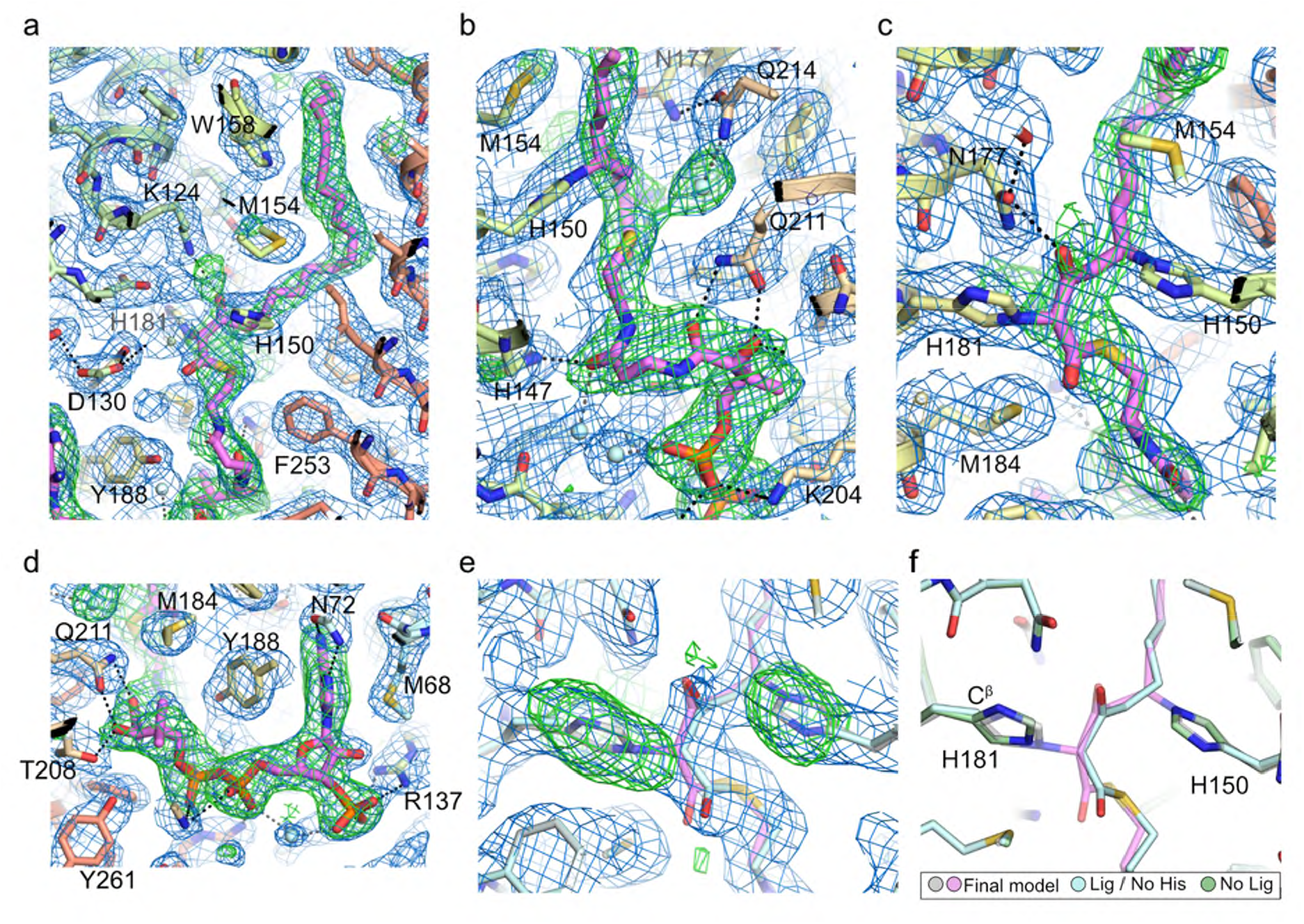
Electron density clearly shows covalently bound 3-keto-eicosanoyl CoA. **a-d**, Electron density running along the catalytic tunnel. Final BUSTER 2mFo-DFc (blue mesh, contoured at 1sigma) and omit mFo-DFc (green mesh, contoured at 2.5sigma) electron density maps are overlaid on the final model. **e**, Comparison of a test refinement in which the imidazole groups of H150 and H181 were removed from the model (grey carbon protein atoms / palecyan ligand carbon atoms) and the final model (palecyan protein carbons / violet ligand carbon atoms). The BUSTER 2mFo-DFc (blue mesh, contoured at 1sigma) and mFo-DFc (green mesh, contoured at 3sigma) maps for the refined histidine-truncated model / unlinked acyl-CoA are shown.. **e**, Comparison of various refined models (green carbons - protein only; palecyan - protein without H150/181 sidechain plus ligand; grey/violet - final model with covalently attached ligand).

**Extended Data Fig.4.**
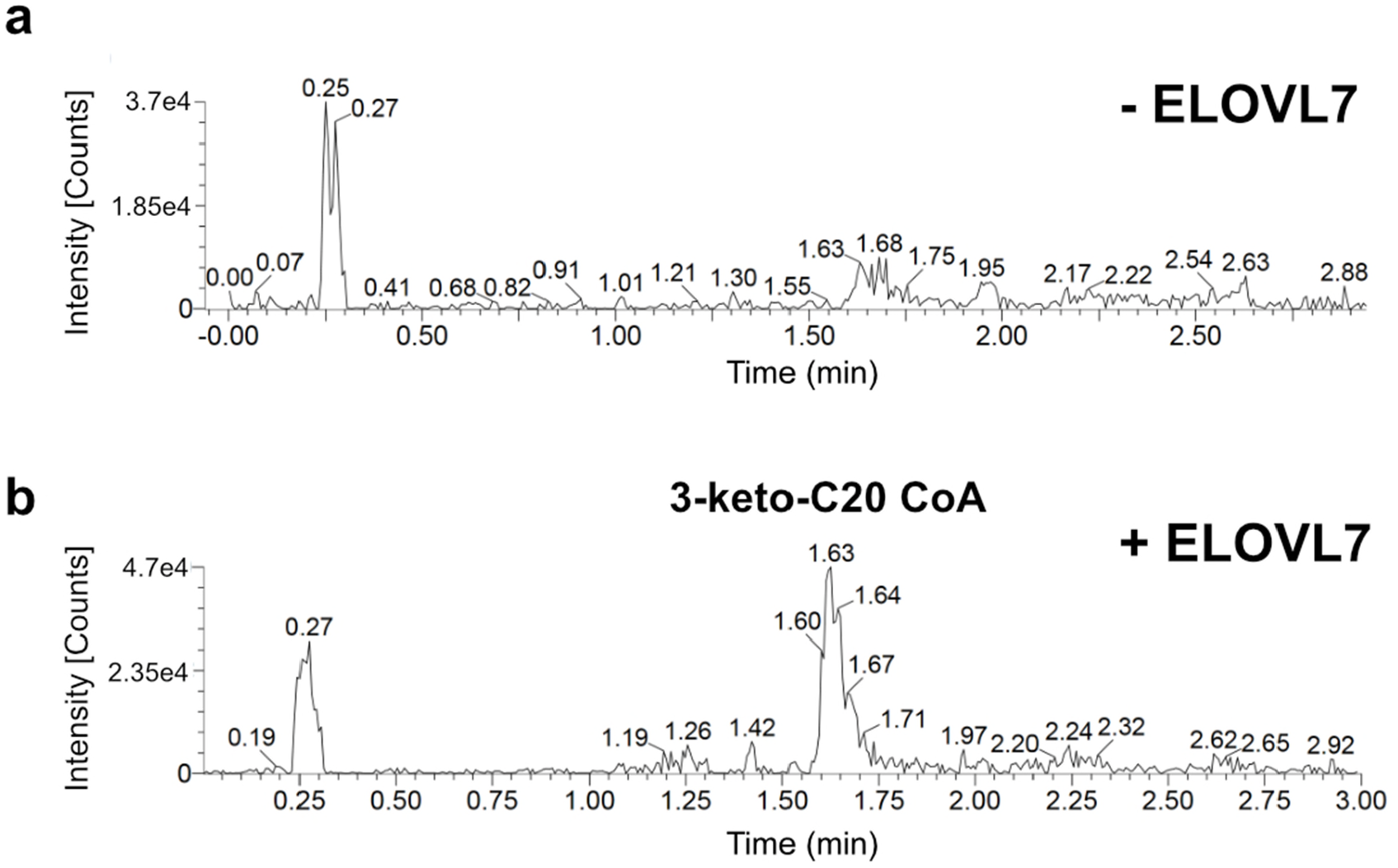
WT ELOVL7 activity. **a**,**b** Activity of residual WT enzyme on incubation with palmitoyl-CoA (C18:0) and malonyl-CoA. Selected ion recording is shown for **a**, reaction mixture without added enzyme and **b**, reaction mixture after 3hr incubation with ELOVL7 enzyme. Ion peak at 1.61 mins corresponds to the expected 3-keto-eicosanoyl (C20)-CoA product of the elongation reaction.

**Extended Data Fig.5.**
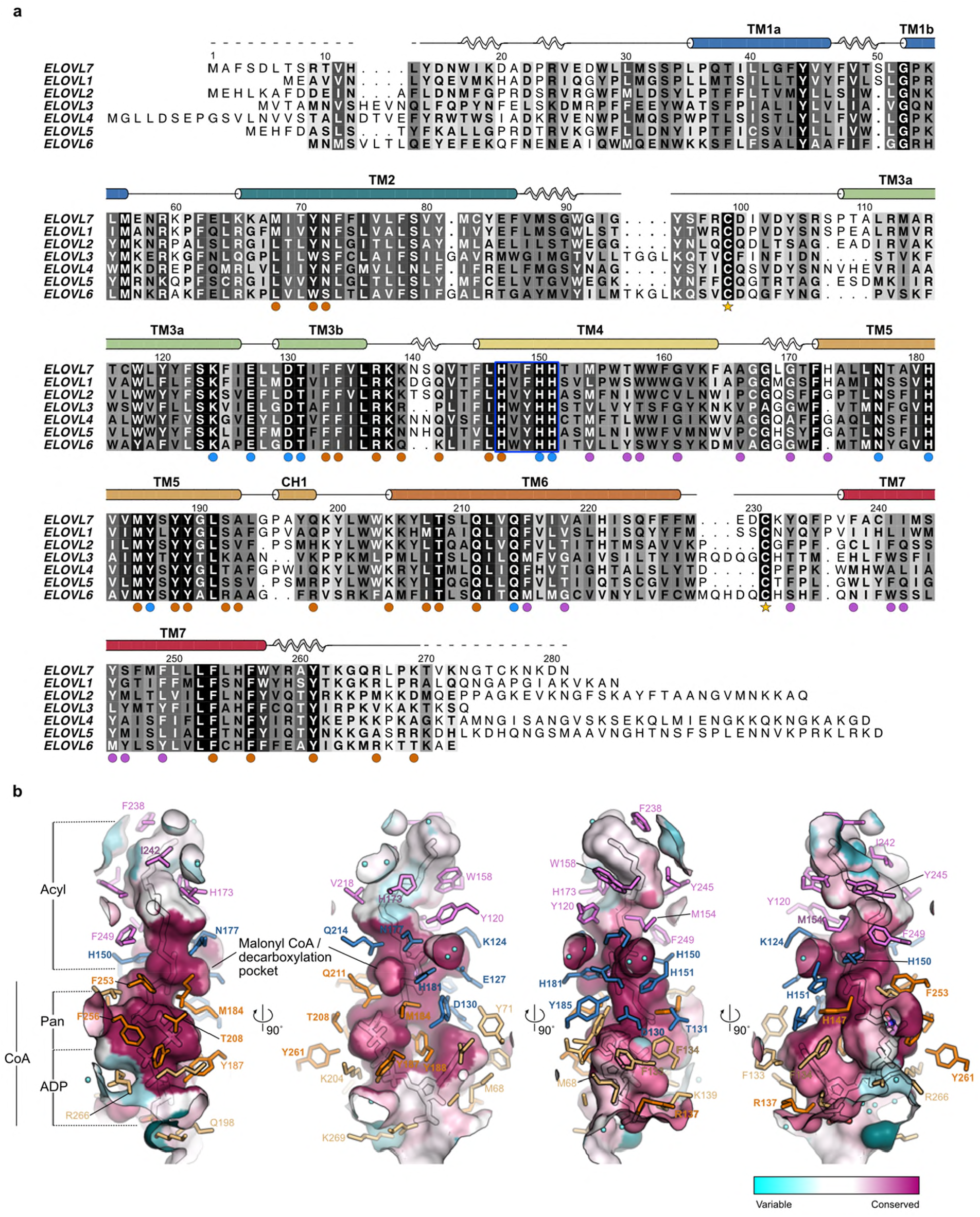
Sequence alignment and active site conservation of human ELOVL family members. **a**, Sequence alignment of human ELOVL1-7. The conserved histidine box (^147^HxxHH^151^) is highlighted by a blue box. Filled circles below alignment indicate residues with a proposed catalytic role (blue) and residues interacting with either the CoA (orange) or acyl (plum) portion of the substrate. Cysteines that form the disulphide bridge (C99-C231) between the TM2/3 and TM6/7 loops are indicated by stars. **b**, Conservation of active site tunnel. Molecular surface representation is coloured by amino acid conservation score calculated by CONSURF analysis^49^ of a diverse set of ELOVL1-7 family members. The various subregions of the tunnel are indicated (ADP / Pan from CoA and Acyl chain). Amino acid residues that form the binding tunnel are coloured according to region (pink, acyl; blue, catalytic site; orange CoA binding).

**Extended Data Fig.6.**
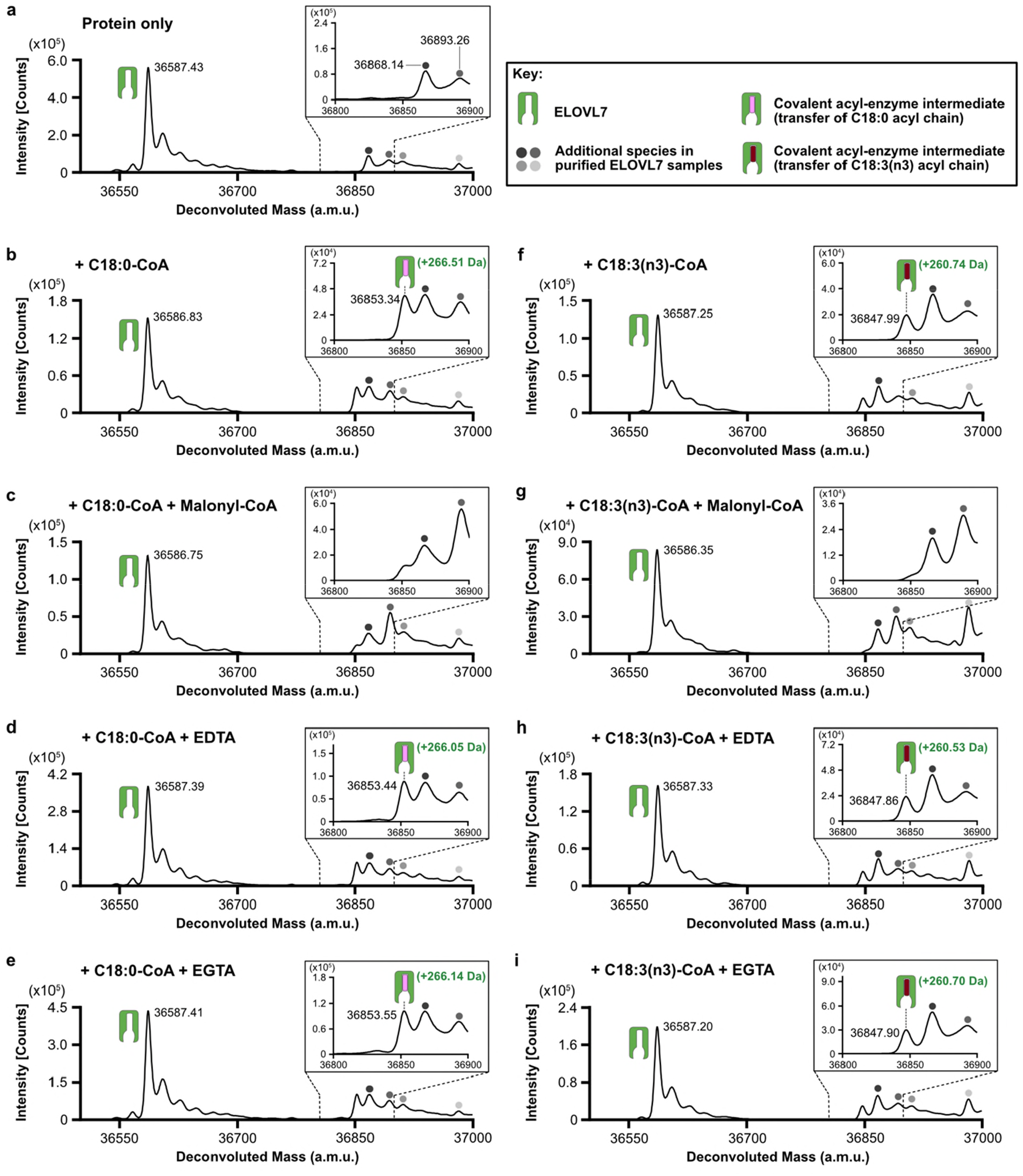
Identification of a covalent acyl-enzyme intermediate of ELOVL7. Purified, tagged, wild-type ELOVL7 was incubated for 2h at 37°C in the presence and absence of known substrates and metal-chelating agents prior to LC-ESI-MS intact mass analysis. Deconvoluted intact mass spectra for ELOVL7 incubated **a**, in the absence of substrates. **b**, with 100μM C18:0-CoA. Expected mass addition for acyl intermediate upon reaction with C18:0-CoA: +266.47 Da. **c**, Incubation of ELOVL7 with 100μM C18:0-CoA and 100μM malonyl-CoA prevents covalent acyl-enzyme intermediate accumulation. **d-e**, ELOVL7 incubated with 100μM C18:0-CoA in the presence of **d**, 1mM EDTA or **e**, 1mM EGTA, showing that intermediate formation is not inhibited by metal-chelating agents. **f**, ELOVL7 incubated with 100μM C18:3(n3)-CoA. Expected mass addition for acyl intermediate upon reaction with C18:3(n3)-CoA: +260.42 Da. **g**, Incubation of ELOVL7 with 100μM C18:3(n3)-CoA and 100μM malonyl-CoA prevents covalent acyl-enzyme intermediate accumulation. **h-i**, ELOVL7 incubated with 100μM C18:3(n3)-CoA in the presence of **h**, 1mM EDTA or **i**, 1mM EGTA. All experiments were repeated independently twice with similar results (n=2 biological repeats, see Supplementary Information).

**Extended Data Fig.7.**
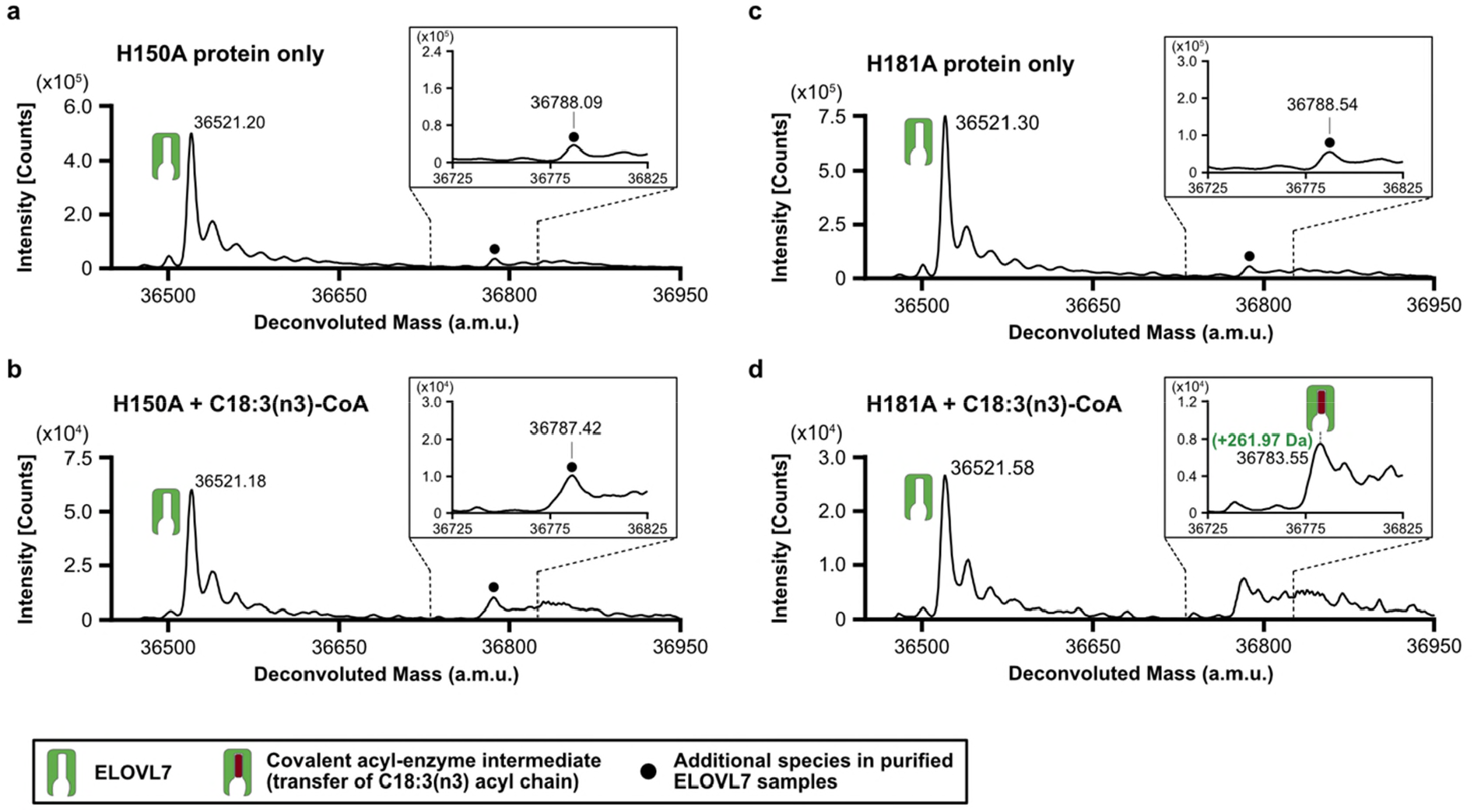
Covalent acyl-enzyme intermediate is formed upon substrate reaction at His150. LC-ESI-MS intact mass analysis of mutant proteins upon incubation at 37°C for 2h. **a-b**, His150Ala mutant protein incubated **a**, in the absence of substrate or **b**, in the presence of 100 μM C18:3(n3) CoA. No acyl-enzyme intermediate formation could be detected with the His150Ala mutant protein. **c-d**, His181Ala mutant protein incubated **c**, in the absence of substrate or **d**, in the presence of 100μM C18:3(n3) CoA. Expected mass shift upon reaction with C18:3(n3) CoA: +260.42 Da. The presence of a background peak at ∼36788 Da precluded testing with C18:0 CoA. All experiments were repeated twice with similar results (n=2 biological repeats, see Supplementary Information).

**Extended Data Fig.8.**
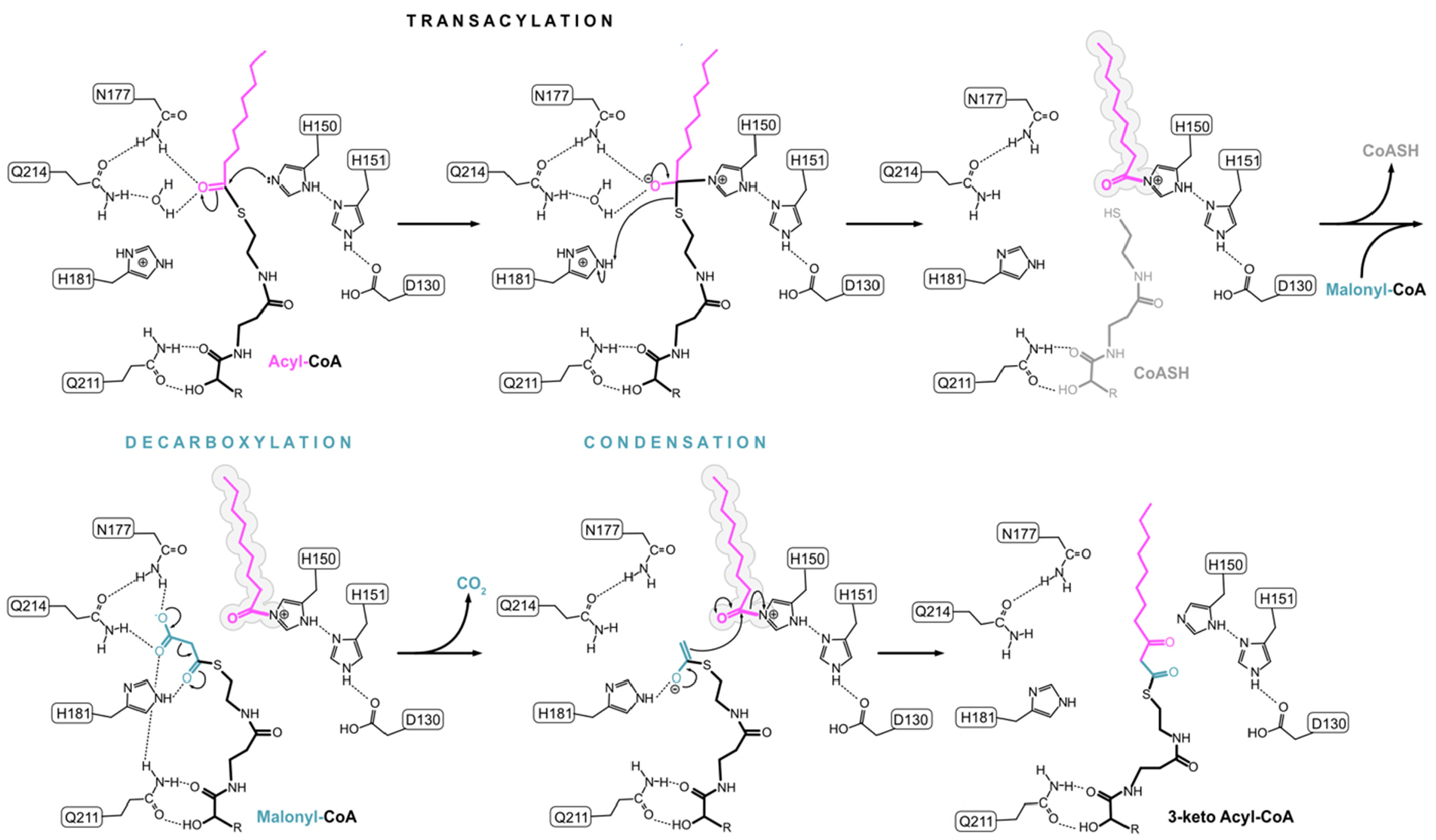
Proposed ping-pong reaction mechanism for ELOVL7.

**Extended Data Table 1.**
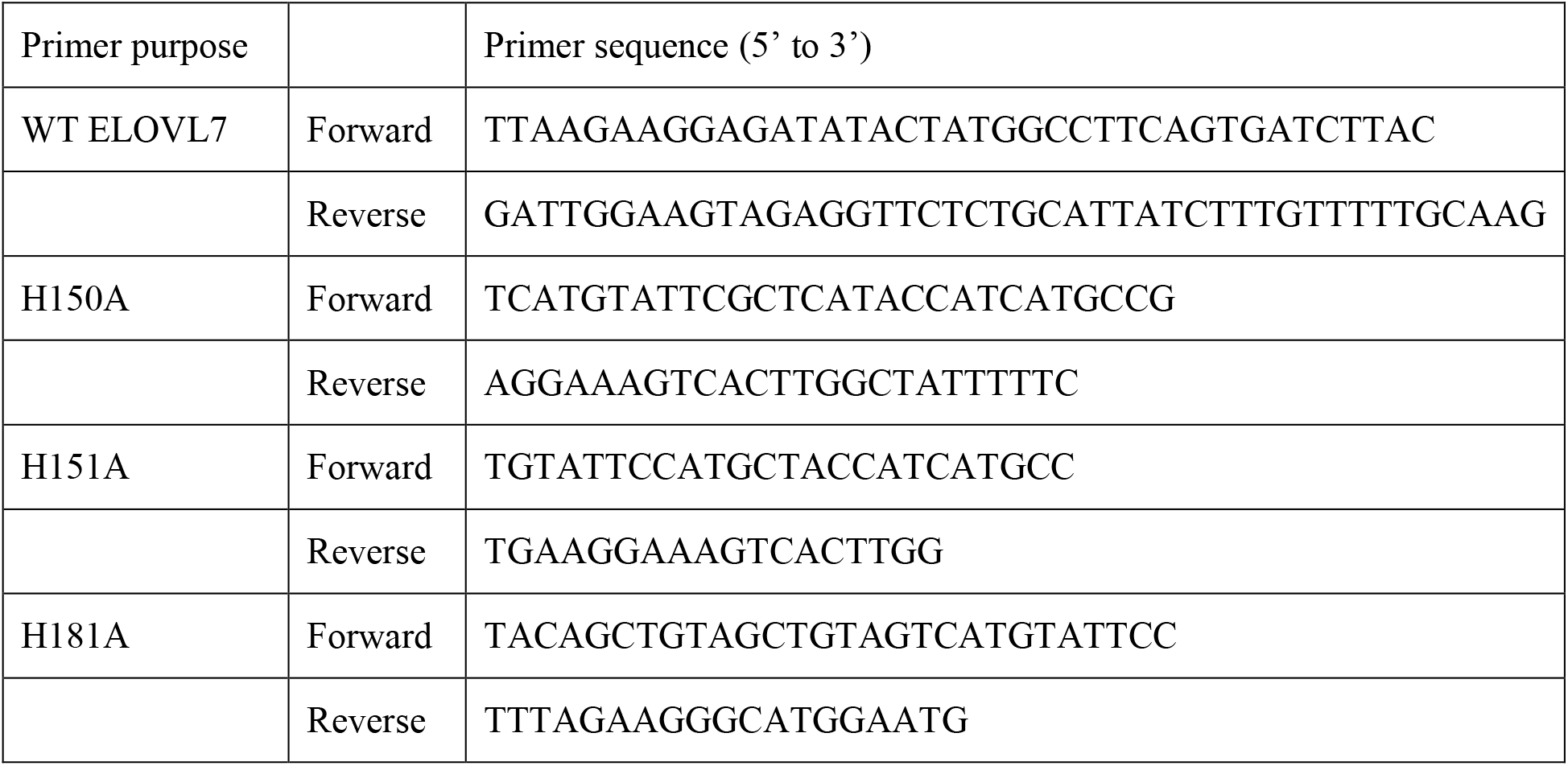
Mutagenesis primer sequences.

## Notes

### Competing Interest Statement

The authors have declared no competing interest.

